# Neuroendocrine influences on dynamic cerebrovascular function and implications for functional MRI

**DOI:** 10.64898/2026.06.06.729907

**Authors:** Melissa E. Wright, Ian D Driver, Cassandra Crofts, Saajan Davies, Jessica J. Steventon, D Samuel Schwarzkopf, Kevin Murphy

## Abstract

To fully profile how ovarian hormones interact with cerebrovascular function, it is vital to consider, not just resting physiology, but also dynamic aspects of cerebrovasculature that support neural activity. This study uses hypercapnic cerebrovascular reactivity (CVR) and the visually-evoked haemodynamic response function (HRF) to investigate the influence of menstrual-related changes in oestradiol and progesterone on dynamic aspects of the cerebrovascular system.

20 menstruating females (age mean[SD]=23.01[4.01]years) completed a 3T MRI scanning session during the early follicular, late follicular, and mid-luteal phases of their menstrual cycle. Circulating hormones were measured via blood samples. Simultaneous blood oxygen level dependant (BOLD)-CVR and cerebral blood flow (CBF)-CVR data were collected using a pseudocontinuous arterial spin labelling (pCASL) acquisition using a dual-excitation (DEXI) readout during periods of hypercapnia (5% CO_2_). The HRF was estimated using a whole brain EPI scan during high-contrast radial checkerboard presentation.

Both oestradiol and additional progesterone variance were associated with increased CVR (both BOLD-CVR and CBF-CVR; p<0.001) and altered HRF shape (p<0.005). No statistically significant regional effects were found.

A secondary experiment investigated the impact of using either canonical or individually mapped HRF in a standard fMRI processing pipeline; namely, population receptive field (pRF) mapping. Results across phases suggest that neither hormone was associated with pRF size when modelled using a canonical HRF (both p>0.05). However, a significant neuroendocrine influence on pRF sizes was discovered when using individually measured HRFs (p<0.001).

This study found evidence that dynamic cerebrovascular functions are sensitive to menstrual-related ovarian hormones, which may be a potential mechanism underlying menstrual symptomatology and has implications for fMRI studies that assume intact neurovascular coupling processes in women, regardless of menstrual staging, to make inferences about neural activity.

## 1. Introduction

Ovarian hormones, notably oestrogen and progesterone, play an important role in healthy cerebrovascular function. Animal model research demonstrates a marked vasodilatory influence of these hormones (Robison et al., 2019; Tostes et al., 2003), which is mirrored in elevated resting perfusion with increasing hormones in human endocrine states such as the menstrual cycle (Cote et al., 2021; Wright et al., 2026). Furthermore, menopause, characterised by a dysregulation and decline in ovarian hormones, is linked to a decrease in vascular health (Hayward et al., 2000; Uddenberg et al., 2024; Wood Alexander et al., 2024), which is associated with cognitive decline (Wood Alexander et al., 2024). Fully profiling and understanding this relationship is vital to highlight periods of cerebrovascular risk across a woman’s lifespan and possible mechanistic pathways for the beneficial role of hormones. However, this remains under-researched and the majority of human neuroimaging research into this topic has focused on baseline vascular function, such as resting cerebral blood flow (CBF), arterial arrival time, and vessel density (e.g., Cote et al., 2021; Guo et al., 2021; Wright et al., 2026). To fully profile how these hormones interact with cerebrovasculature, it is essential to also consider dynamic vascular physiology that supports neural function.

The cerebrovascular system not only delivers a sufficient level of oxygenated blood at baseline but also adjusts local blood flow in response to tissue demands. This dynamic aspect of cerebrovascular physiology is a key indicator of healthy cerebrovascular reserve capacity, and is essential to meet the increased energy demand following neural activation. This dynamic aspect can be assessed via cerebrovascular reactivity (CVR), the ability of blood vessels to dilate in response to a vasoactive stimulus, such as CO_2_, and thus increase local CBF. Impairments have been associated with cognitive decline (Catchlove et al., 2018; Liu et al., 2024), neurodegenerative diseases (Aslanyan et al., 2024; Chen, 2018; Marshall et al., 2014; Ryman et al., 2023; Tang et al., 2025) and wider vascular health (Hobden et al., 2025). Increased vascular reactivity in the carotid and cerebral arteries has been reported with menstrual-related increases in ovarian hormones (Debert et al., 2012; Krejza et al., 2013; Skinner et al., 2023), but this has not yet been comprehensively investigated across the cortex using high-resolution MRI.

The characterisation of the haemodynamic response function (HRF) is another useful metric to interrogate the dynamic functionality of the vascular system. Using blood oxygen level dependant (BOLD) functional MRI (fMRI), the BOLD signal is mapped in response to a stimulus designed to evoke a strong but brief neural response. The final HRF will contain information about both the neural activation, cerebrovascular coupling mechanisms, and local blood flow increases. While it is not possible to detangle specific contributions, it provides a useful overview of the whole process. The HRF shape is also essential for many fMRI analysis pipelines (such as retinotopic population receptive field (pRF) mapping of the visual cortex) which rely on the assumption that there are no systematic changes to the HRF in order to make assumptions about the underlying neural activity. Multiple studies have reported menstrual-dependant changes in neural activation based on the BOLD fMRI signal (Alonso-Alonso et al., 2011; Zhu et al., 2010); to have greater confidence in inferences from these results, it is essential to map the HRF itself throughout the menstrual cycle.

This study therefore aims to utilise hypercapnic CVR and visually-evoked HRFs to investigate the contribution of menstrual-related changes in oestradiol (the most common form of menstrual oestrogen) and progesterone to these dynamic aspects of the cerebrovascular system. It is hypothesised that, in line with previous Doppler imaging studies (Debert et al., 2012; Krejza et al., 2013; Skinner et al., 2023), we will find an association between circulating hormone level and increased CVR and HRF peak amplitude. As a secondary experiment, the influence of using either a canon or individualised HRF on a standard fMRI processing pipeline will then be investigated; if there is a neuroendocrine influence on HRF shape, this may bias the outcome of fMRI results that utilise the HRF for signal convolution.

## 2. Experiment 1: Establishing endocrine influences on dynamic vascular functions

### 2.1. Methods

MRI scanning was completed three times across a menstrual cycle (in the early follicular phase [EFP; cycle day 1-4], late follicular phase [LFP; cycle day 10-12], and mid-luteal phase [MLP; cycle day 20-22]), according to the self-reported onset day of menses. Participants were required to fast for a minimum of 4 hours and abstain from caffeine, alcohol, and strenuous physical activity for 12 hours before a test session. All sessions took place at a similar time of day.

#### 2.1.1. Participants

The sample described here were recruited as part of a previous study on baseline cerebral and retinal vascular physiology across the menstrual cycle (Wright et al., 2026). Twenty regularly-menstruating participants (age mean[SD]=23.01[4.01] years; visual acuity mean[SD]=-0.21[0.22] logMAR) completed an initial screening session to determine eligibility and to give them the opportunity to ask any questions about the study (EFP=17; LFP=15; MLP=18). During the first session they gave written informed consent. Eligible participants had a regular (i.e., self-reported cycle length between 27-31 days for at least the last three cycles) menstrual cycle. Participants were excluded if they had any commonly accepted contraindications to MRI scanning, a clinically significant condition (such as diabetes or a neurological, cardiovascular, psychiatric, cerebrovascular, or respiratory condition), were clinically vulnerable or extremely vulnerable to COVID-19, were pregnant or had been in the last 6 weeks, took hormonal contraceptives presently or in the last 6 months, or were post-menopausal. A COVID-19 screening questionnaire was also required before each session and testing sessions were rearranged if participants reported any COVID-related symptoms. Participants received financial compensation for each attended testing session. Ethical approval was given by the Cardiff University School of Medicine Research Ethics Committee and was in accordance with the Declaration of Helsinki.

This sample met our calculated sample size threshold for 90% power (n=14 required based on effect size dz=0.98 in ΔMCAv [middle cerebral artery velocity] (Peltonen et al., 2016)), initially calculated for the previous resting study (Wright et al., 2026).

#### 2.1.2. Hormone sampling

Venepuncture was used to sample 5ml of blood from the hand/arm. The anonymised sample was sent to the Medical Biochemistry Laboratories at the University Hospital of Wales for measurement of oestradiol and progesterone serum concentration with NHS standard protocols. Oestradiol levels were measured with an Abbott Architect analyser and progesterone levels were measured with an Abbott Alinity analyser.

As was done previously (Wright et al., 2026), progesterone values were transformed to reduce the influence of intercorrelation on the statistical models. Oestradiol levels were regressed from progesterone levels, and the model residuals were taken as the variance of progesterone that is independent from oestradiol, hereby referred to as ‘resProgesterone’.

#### 2.1.3. MRI session – acquisition

Data were acquired at Cardiff University Brain Research Imaging Centre (CUBRIC) on a Siemens MAGNETOM Prisma 3T scanner with a 32-channel head coil.

- *Cerebrovascular reactivity [CVR]* – Simultaneous BOLD and CBF reactivity data was collected using a pseudocontinuous arterial spin labelling (pCASL) acquisition using pre-saturation and background suppression (Okell et al., 2013) and a dual-excitation (DEXI) readout (Germuska et al., 2019; Schmithorst et al., 2014). The parameters include: maximum repetition time [TR]=3600ms, echo time [TE]1=10ms, TE2=30ms, in-plane resolution=3.4×3.4mm, slice thickness=6.5mm, GRAPPA acceleration factor=3, slices=15. The post label delay (PLD) and tag duration were both 1500ms. The acquisition time for this sequence was 9 minutes 11 seconds and includes interleaved periods of hypercapnic and medical air gas delivery (detailed below). A calibration (M0) image was acquired for CBF quantification and background suppression switched off (TR=6s, TE=10ms, both AP and PA readout).
- *Haemodynamic response function [HRF]*– This was undertaken with a whole brain EPI, while a checkerboard stimulus was presented (detailed below). Parameters include: TR=2s, TE=30ms, resolution=2mm^3^, flip angle=58°, 64 slices, FOV=220mm, multi-band acceleration factor=4. To aid in distortion correction, phase and magnitude B0 mapping images were also taken.
- *Structure* – A high-resolution structural T1 image was collected via a Magnetization-Prepared Rapid Gradient-Echo (MPRAGE) scan. Parameters include: TR=2.1s; TE=3.24ms; inversion time [TI]=850ms, resolution=1mm^3^.

#### 2.1.4. MRI gas challenge

During acquisition of the pCASL sequence, interleaved periods of either hypercapnic (5% carbon dioxide [CO_2_], balance air) or normocapnic gas mixture was delivered to participants via a facemask, using a block design. To monitor and measure physiological data during the scan, a pulse oximeter (Biopac Systems Inc, California, USA), facemask, and custom-built respiratory belt were secured. Partial pressures of end-tidal O_2_ (PETO2) and CO_2_ (PETCO2) were measured from expired gases sampled from the participant’s facemask using an AD Instruments gas analyser and data sampling system (PowerLabVR, ADInstruments, Sydney, Australia). Due to either technical difficulties or participants opting out of the gas challenge, a reduced sample size was available for the CVR analysis (Total N: 19; EFP=15; LFP=9; MLP=13).

#### 2.1.5. fMRI stimuli

MRI compatible vision correction was supplied if needed and all participants confirmed that the whole stimulus screen was in focus. The stimulus screen size was 25°x14° at a viewing distance of ∼88cm.

The HRF was mapped with a high-contrast black-and-white alternating radial checkerboard (see Figure 1). The radial component comprised of concentric bands with an average spatial frequency of approximately ∼1.3 cycles per degree along the eccentricity axis (decreasing with increasing eccentricity). This was presented to participants for 2 seconds (1 TR), followed by a blank grey screen for 20 seconds (10 TR). This sequence was repeated ten times while the participant was asked to maintain focus on a central fixation. Twenty seconds of mean luminance was also included at the start of the sequence to establish a baseline of no stimulation, which was then removed for further processing and analysis.

**Figure 1.**
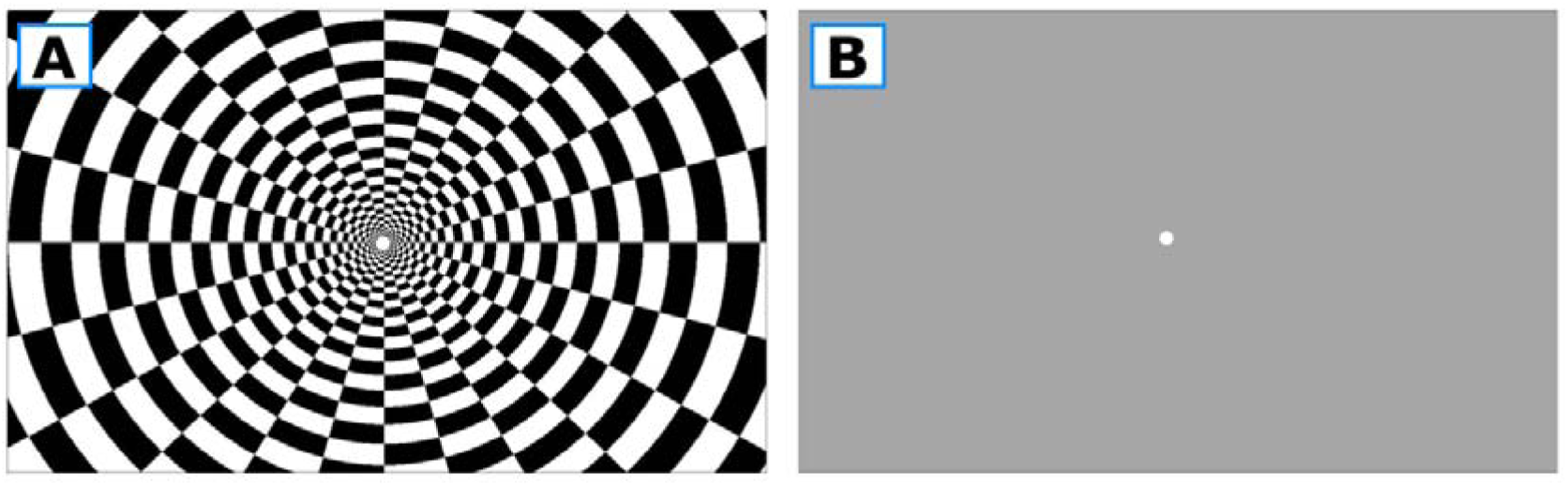
The stimulus used for Haemodynamic Response Function (HRF) mapping in the scanner. A: The radial checkerboard stimulation. B: Mean grey rest stimuli. Checkerboard stimulation (A) is displayed for one Repetition Time (TR; 2 seconds), followed by ten TR of rest stimuli (B; 20 seconds). This was repeated ten times.

#### 2.1.6. MRI pre-processing

*Structural grey matter mask –* A grey matter mask was generated to be used in other processing pipelines. The T1 MPRAGE structural images were processed using the fsl_anat pipeline (Jenkinson et al., 2012; Smith et al., 2004; Woolrich et al., 2009), which includes reorientating to standard orientation, bias-field correction, brain extraction, and tissue-type segmentation (FAST; Zhang et al., 2001).

### Cerebrovascular reactivity

The dual-echo CVR data was first split into ASL and BOLD data. Pre-processing of the separate timeseries (i.e., motion correction, slice time correction, conversion to % BOLD signal change) was completed using the Analysis of Functional NeuroImages (AFNI) software package (Cox, 1996; provided in the public domain by the National Institutes of Health, Bethesda, MA, USA; http://afni.nimh.nih.gov/afni). To quantify CBF from the ASL data, several additional processing steps were taken. Firstly, the deltaM (i.e., the tag minus control signal change in magnetisation) was calculated. The M_0_ of blood was calculated as:

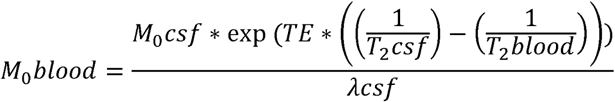

λ*csf* = blood-CSF partition coefficient (1.15; Herscovitch & Raichle, 1985; Pinto et al., 2020)

*M*_0_*csf* = the M_0_ of the cerebral spinal fluid (CSF) was taken from a CSF mask generated around a manually positioned central point in the lateral ventricles.

*TE* = echo time (0.01s)

*T*_2_*csf* = T_2_* of CSF (0.4; Pinto et al., 2020)

*T*_2_*blood* = T_2_* of blood (0.06s; Zhao et al., 2007)

A non-linear fitting algorithm (AFNI’s 3dNLfim function) and the general kinetic perfusion model provided final CBF quantification (Buxton et al., 1998). Arterial arrival time was estimated at 0.7s (based on the average AAT in grey matter for women reported in MacIntosh et al., 2010).

PETCO_2_ traces recorded during the gas challenge were processed within MATLAB software (The MathWorks Inc., 2021), using in-house scripts. These scripts rescale the PetCO_2_ to mmHg and convert the trace to a HRF-convolved regressor. This final regressor is then cross-correlated to the global BOLD or ASL time course (with 41 incremental time shifts in PetCO_2_ trace) to find the best temporal match between both timeseries, defined as the correlation with the highest rho value.

CVR maps were then generated as either BOLD signal percent change from baseline or CBF ml/100g/min per unit of PETCO_2_ by convolving the MRI timeseries with the processed PETCO_2_ trace using the 3dDeconvolve AFNI command. A model fit threshold of R^2^>0.1 was applied to the final CVR maps. A transformation matrix was generated using FLIRT (Jenkinson et al., 2012) to register the Desikan-Killiany Atlas to each participants’ native space. The cortical regions were further restricted to the individual’s grey matter. The median values were extracted for each region (Desikan et al., 2006). The average CVR across all surviving voxels were taken as global measures for comparison.

### Haemodynamic response function mapping

EPI data underwent B0 distortion correction using the FMRIB Software Library (FSL) v.6. software package (Jenkinson *et al*., 2012). The rest of the pre-processing steps were completed using the Analysis of Functional NeuroImages (AFNI) software package (Cox, 1996; provided in the public domain by the National Institutes of Health, Bethesda, MA, USA; http://afni.nimh.nih.gov/afni). This included motion correction, 5mm spatial smoothing and scaling the timeseries so the output represents percentage signal change. The timeseries was averaged across presentations and a gamma variate HRF model fit using the 3dDeconvolve and 3dNLfim commands.

The data was then extracted from a series of anatomical regions from the Desikan-Killiany Atlas (restricted to the grey matter mask, as above), which were all assumed to be responsive to visual stimuli. The regions of interest were as following (right and left hemisphere):

- Pericalcarine sulcus
- Lateral occipital cortex
- Cuneus
- Fusiform gyrus
- Lingual gyrus

In order to describe the full HRF shape, data was interpolated using a cubic convolution in MATLAB 2021a (The MathWorks Inc., 2021). Three metrics were extracted from these fits to describe the final HRF shape and allow for quantitative comparisons between groups. These metrics were the Peak Amplitude (PA), the Time-To-Peak (TTP, defined as the corresponding time [in seconds] to reach the PA), and the Full-Width-Half-Maximum (FWHM, measured across the initial positive Gaussian peak in seconds). Any negative-going HRFs were excluded (left lateral occipital cortex=2/49; right lateral occipital cortex=3/49, left cuneus=22/49, right cuneus=25/49, left fusiform=8/49, right fusiform=6/49, left lingual=10/49, right lingual=8/49, left pericalcarine=10/49, right pericalcarine=9/49). For each participant, all surviving ROIs were averaged to create a global comparison value.

#### 2.1.7. Statistical Analysis

The contribution of circulating hormone levels to the variance in the outcome variables was calculated using linear mixed models (LMMs). Statistical analysis was completed in R software (v.4.2.2; R core team, 2022) and LMMs were constructed using the ‘lmerTest’ analysis package (Kuznetsova et al., 2017). Statistical significance was set at the P<0.05 level and p-values were generated by statistically comparing the model with and without the outcome of interest (ANOVA), which indicates whether that outcome significantly contributes to model variance. With all LMMs, residual plots were visually inspected to ensure no notable deviations from homoscedasticity or normality. If residual plots notably deviated, a robust mixed-effect model (Huber weighting) was used instead, which downweighs the influence of residual outliers. For robust models, p-values were calculated using robust t-values and Satterthwaite approximations of degrees-of-freedom, as has been done previously (Geniole et al., 2019). In all models, oestradiol, resProgesterone, and ROI (if applicable) were used as fixed effects, with participant as a random effect. An interaction term between ROI and hormone levels was then added to investigate if the effect is driven by a particular ROI (if applicable).

#### 2.1.8. Sensitivity analysis

Sensitivity analyses were also carried out to investigate whether any CVR effects could be attributed to other effects, such as task-related effort and ventilatory drive. The CVR quality check metrics were:

- ΔPETCO_2_—defined as the difference between the normocapnia and hypercapnia condition end tidal CO_2_ recordings.
- CO_2_-BOLD Rho—the Rho statistic from the Spearman’s Rho correlation between the optimum PETCO_2_ trace and the BOLD time course, indicating the direction and strength of the relationship.
- Optimal CO_2_-BOLD lag—the time lag that the PETCO_2_ trace was shifted to get the optimal correlation with the BOLD time course (defined as the highest Rho value).

Robust LMMs were used to investigate whether ovarian hormones had a significant association with any of these factors, similar to those LMMs used for the main outcomes.

### 2.2. Results

#### 2.2.1. Vascular outcomes

Statistics are presented in Tables 1-2. As the residuals from both BOLD- and CBF-CVR were found to deviate from normality, robust models were used.

**Table 1.** Results from Robust Models for CVR-based outcomes CI= confidence interval; CVR= cerebrovascular reactivity; CBF= cerebral blood flow; BOLD= Blood oxygen level dependant.

|  | **Outcome** | **t-**<br>**statistic** | **p-value** | **Estimate per**<br>**unit increase** | **Upper and**<br>**lower 95% CI** |
| --- | --- | --- | --- | --- | --- |
| **CBF-CVR** | *Oestradiol* | 8.003 | <0.001 | 0.0008 | 0.0006-0.0010 |
|  | *resProgesterone* | 11.294 | <0.001 | 0.0208 | 0.0172-0.0244 |
| **BOLD-**<br>**CVR** | *Oestradiol* | 11.072 | <0.001 | 0.0001 | 0.00008-0.00012 |
|  | *resProgesterone* | 4.193 | <0.001 | 0.0007 | 0.0004-0.0011 |

**Table 2.** Results from Linear Mixed Models for HRF-based outcomes Df=1. Chi-squared and p-values are generated by statistically comparing the model with and without the outcome of interest (ANOVA), which suggests whether that outcome statistically significantly contributes to model variance. CI= confidence interval.

|  | **Outcome** | **$\chi^2$** | **p-value** | **Estimate per**<br>**unit increase** | **Upper and**<br>**lower 95% CI** |
| --- | --- | --- | --- | --- | --- |
| **HRF – PA** | *Oestradiol* | 21.20 | <0.001 | 0.0002 | 0.0001-0.0003 |
|  | *resProgesterone* | 0.08 | 0.77 |  |  |
| **HRF – TTP** | *Oestradiol* | 9.44 | 0.002 | -0.0009 | -0.0014-<br>-0.0003 |
|  | *resProgesterone* | 0.19 | 0.66 |  |  |
| **HRF –**<br>**FWHM** | *Oestradiol* | 2.99 | 0.08 |  |  |
|  | *resProgesterone* | 24.02 | <0.001 | 0.025 | 0.0153 – 0.0352 |

Oestradiol was found to explain a significant amount of global CBF-CVR variance (*t-statistic*=8.00; p<0.001; estimate[95% CIs]=0.0008[0.0006-0.0010]). While the CBF perfusion level would increase by 2.695ml/100g/min per 1mmHgCO_2_ in a ‘zero hormone’ condition therefore, this fixed effect estimate suggests that in a high oestradiol environment (e.g., 508pmol/L, the average increase in this sample), CBF perfusion level would increase by 3.101ml/100g/min per 1mmHgCO_2_. A significant effect was also found with resProgesterone and CBF-CVR (*t-statistic*=11.29; p<0.001; estimate[95% CIs]=0.02[0.0172-0.0244]). For a condition with high levels of resProgesterone (20nmol/L, the average increase in this sample), CBF would increase by 3.095ml/100g/min per 1mmHgCO_2_. This is placed in the context of a 5% CO_2_ increase in Figure 2.

**Figure 2.**
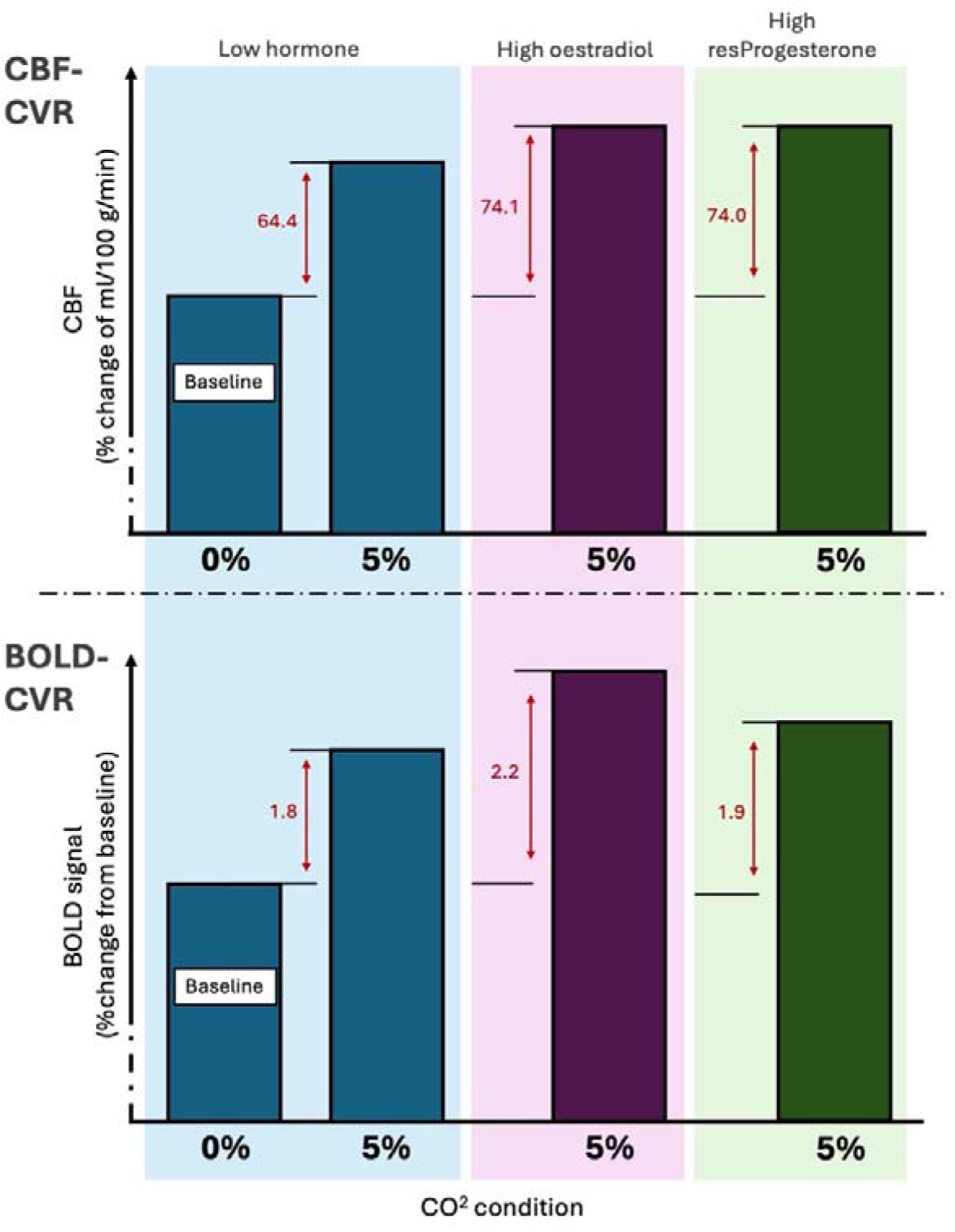
Illustration of cerebrovascular reactivity (CVR) changes across hormone and CO2 conditions. 5% CO2 is taken as 7.5mmHg, based on the average difference between the normocapnia and hypercapnia condition end tidal CO2 recordings in this cohort. ‘High’ oestradiol and high resProgesterone are taken as 508pmol/L and 20nmol/L respectively, based on average increases measured across the menstrual cycle in this sample. Baseline CBF values are based on model intercepts. CBF= cerebral blood flow; BOLD= Blood oxygen level dependant.

Significant associations were also observed for BOLD-CVR. Both oestradiol (*t-statistic*=11.07; p<0.001; estimate[95% CIs]=9.93×10^-5^[8.17×10^-5^-1.17×10^-4^]) and resProgesterone (*t-statistic*=4.19; p<0.001; estimate[95% CIs]=7.28×10^-4^[0.0004-0.0011]) were associated with a small but significant increase in BOLD-CVR (see Figure 2).

In terms of HRF shape, increasing oestradiol was associated with slightly higher PA (*χ*^2^(1)=21.20; p<0.001; fixed effect estimate=1.94×10^-4^%BOLD; 95% CIs=0.0001-0.0003; average variation attributed to oestradiol across a cycle in this sample=0.10%BOLD) and earlier TTP (*χ*^2^(1)=9.44; p=0.002; fixed effect estimate=-8.54×10^-4^ seconds; average variation attributed to oestradiol across a cycle in this sample=0.43 seconds). Increasing resProgesterone was associated with wider FWHM (*χ*^2^(1)=24.02; p<0.001; fixed effect estimate=0.025 seconds; 95% CIs=0.015–0.035; average variation attributed to resProgesterone across a cycle in this sample=0.51 seconds). Effect illustrated in Figure 3.

**Figure 3.**
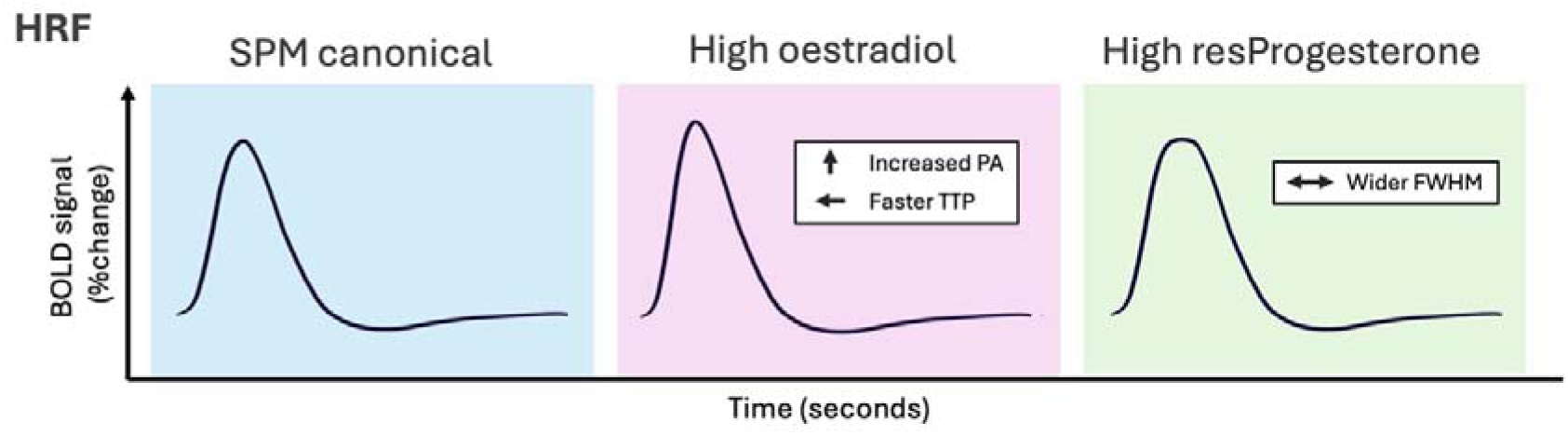
Schematic illustration of HRF changes across hormone levels. ‘High’ oestradiol and high resProgesterone are taken as 508pmol/L and 20nmol/L respectively, based on average increases measured across the menstrual cycle in this sample. SPM canonical HRF shape used to demonstrate baseline ‘no hormone’ level.

No ROI analyses were statistically significant across CVR and HRF metrics, suggesting the significant results above were global effects.

#### 2.2.2. Sensitivity analysis

Neither oestradiol nor resProgesterone explained a statistically significant amount of variance in any of the CVR quality metrics (all P>0.05), suggesting that endocrine effects on these respiratory variables do not explain our results.

## 3. Experiment 2: Implications for fMRI analysis

### 3.1. Rationale

Systematic endocrine-related influence on HRF shape, such as those shown above, may have implications for fMRI analysis, particularly those that use a standardised HRF-convolution of a predicted BOLD signal or event-related design. As a secondary experiment, we therefore aimed to investigate the impact of using either a standard canonical HRF or the individually measured HRF on an fMRI retinotopic pipeline. Specifically, population receptive field (pRF) mapping of the visual field was completed, which uses retinotopic stimuli to assess voxel-level estimates of receptive field sizes (Dumoulin & Wandell, 2008). This pipeline involves HRF convolution of a predicted time course to estimate pRF size and can be particularly sensitive to changes in full-width-half-maximum (FWHM) of the HRF. If associations with endocrine levels were found with the canonical HRF but not the individual HRF, this would suggest that the associations are driven by dynamic vascular functions, while the opposite would imply that neural changes across endocrine states can be masked if vascular changes are not considered. However, it may be that any endocrine influence on the HRF shape is too subtle to be impactful in a standard BOLD fMRI sequence.

### 3.2. Methods

#### 3.2.1. MRI session – acquisition

During all MRI sessions described above, all participants but one (EFP=1; LFP=1; MLP=1) also completed an additional BOLD fMRI scan for pRF mapping (final pRF sample: EFP=16; LFP=14; MLP=17). This was undertaken with a whole brain EPI, while retinotopic stimuli was presented. Parameters include: TR=1s, TE=30ms, resolution=2mm^3^, flip angle=58°, 64 slices, FOV=220mm, multi-band acceleration factor=4. The acquisition time was 2minutes 54seconds and each participant completed this scan 3 times (for later averaging).

The structural MPRAGE and field mapping sequences described above were also utilised in this experiment. Additionally, the HRF EPI scan was re-used for this experiment to generate a surface-based HRF appropriate for this fMRI analysis.

#### 3.2.2. fMRI stimuli

The stimuli used for retinotopic mapping was a moving segment filled with a natural scene, which moved across the screen in a set order. The bar stepped through 18 consecutive positions to cover a circular area of 14° diameter (each bar was 1.48° wide). The step size was half the bar width. Five background scenes were presented at each position for 0.6 seconds each, with three seconds between each position change. The stimuli crossed the screen in the following order: left to right, bottom-right to top-left, top to bottom, bottom-left to top-right, and then the reverse. A final blank epoch was included of 5 TR. In order to maintain fixation, a simple fixation task was used. The fixation dot changed red at random irregular intervals throughout the run, and the participant was required to count in their head the number of times this happened.

The HRF mapping stimulus has been described above (see Figure 1). Briefly, a high-contrast black-and-white alternating radial checkerboard was presented for 1 TR, followed by 10 TR of grey mean luminance. This was repeated ten times.

#### 3.2.3. Structural MRI pre-processing and visual area estimation

The subject-specific structural MPRAGE scans were processed and segmented with FreeSurfer v.7.4.0 tools (Dale et al., 1999; Fischl, 2012). This process divided the structural data into vertices, which could be ‘inflated’ to provide a smoothed projection of the cortical surface for use in functional data surface projection and visualisation.

Objective estimations of primary and secondary visual areas (V1 and V2), independent of either retinotopic map, were generated for later pRF size extraction and averaging. These were generated from the Freesurfer cortical surface based on the probabilistic retinotopic maps on Benson *et al.,* (2014), whereby areas are defined by the likelihood of a given coordinate being associated with a given visual area.

#### 3.2.4. Functional MRI pre-processing

Retinotopic and HRF EPI scans underwent preprocessing using the FSL v.6. software package (Jenkinson *et al*., 2012), which included B0 distortion correction, motion correction (MCFLIRT; Jenkinson et al., 2002) and a 100Hz highpass filter cut-off. Additionally, the HRF data was spatially smoothed using a 5mm FWHM kernel. Following this, registration of the partial-volume EPIs (both retinotopy and HRF runs) to structural data took place in SPM12 (Penny et al., 2007). The functional data were then surface projected onto the smoothed structural scan (0.5 cortex sampling step). A functional time series was created for each vertex for all functional scans and the pRF mapping runs were averaged. Finally, linear detrending and z-score normalisation were applied to these time series.

#### 3.2.5. Surface-based haemodynamic response function estimation

Surface-based HRF fitting was completed using SamSrf software tools (v.10.1; https://github.com/samsrf/samsrf). The BOLD signals generated by the ten checkerboard photic bursts were first averaged within each vertex, excluding any outliers (defined as a time series that demonstrated greater than ±1.5 standard deviation from mean). Analysis was also constrained to only visually active regions to avoid noise. This was defined as a vertex with a signal (averaged over the first five scans) that had a clear response (>1 standard error from the mean) to the photic stimulation. A double-gamma function could then be fitted to the averaged response to estimate the individual visually-evoked HRF fit that could be used in fMRI convolution.

#### 3.2.6. Population receptive field mapping

Population receptive field (pRF) mapping was undertaken within SamSrf MATLAB software (v.10.1; https://github.com/samsrf/samsrf), restricted to an occipital lobe region-of-interest defined by anatomical landmarks on the cortical surface reconstruction. The pRF fitting pipeline has been described in detail previously (Altan et al., 2025; Schwarzkopf et al., 2014). Briefly, the predicted neural response at each voxel is estimated using the linear overlap between the 2D Gaussian pRF model and a binary mask of the stimulus, which is then convolved with a HRF curve. The fit is then optimised between the predicted and measured BOLD response, with a coarse-to-fine approach and a noise ceiling threshold of 0.1. Firstly, a coarse (grid search) fit is completed, using heavily smoothed data (Gaussian kernel full height= 8.3mm) and plausible combinations of pRF parameters. The generated predicted BOLD time series at each point in this search grid were then correlated with the observed BOLD time series. The parameters that led to the best correlation were then optimised further in the second fine fitting stage. The final map contains information on pRF size (defined as the sigma of the Gaussian), retinotopic coordinates (*x* and *y*), *β* amplitude, and R^2^ model fit for each vertex.

Average pRF size for V1 and V2 were then extracted. To ensure that only good quality data were included, only data with R^2^>0.09, pRF sizes within 0.5° and 50° and retinotopic co-ordinates within the stimulated area (≤7°) were included.

This pipeline was repeated twice for each participant and each hemisphere. All analysis steps were identical apart from the 2D HRF shape used in pRF modelling; either the SPM12 canonical HRF or the individually mapped HRF was used.

#### 3.2.7. Statistical Analysis

As above, the contribution of circulating hormone levels to the variance in pRF was calculated using LMMs. Statistical significance was set at the P<0.05 level. With all LMMs, residual plots were visually inspected and showed no notable deviations from homoscedasticity or normality. Oestradiol, progesterone, ROI (primary and secondary visual areas) and hemisphere (left and right) were used as fixed effects, with participant as a random effect. This was completed with either the canonical or individually mapped HRF to investigate the impact of custom HRF convolution.

### 3.3. Results

An example of a retinotopic map is presented in Figure 4 and average pRF sizes for canonical and individually mapped HRF convolutions across menstrual phases are shown in Figure 5.

**Figure 4.**
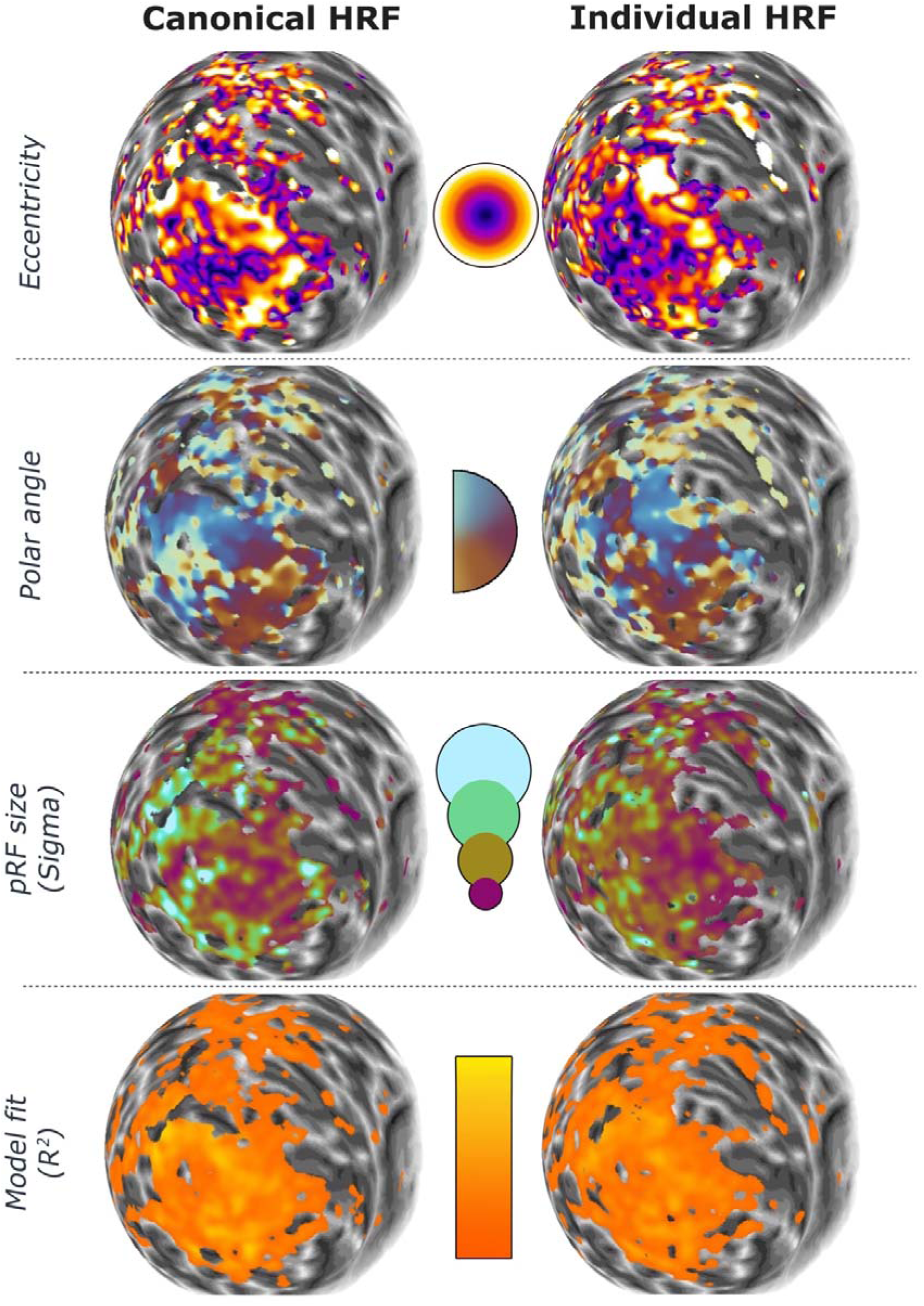
Example retinotopic map for a representative individual. HRF = haemodynamic response function; pRF = population receptive field. Smoothed with spherical FWHM of 5mm to aid with visualisation.

**Figure 5.**
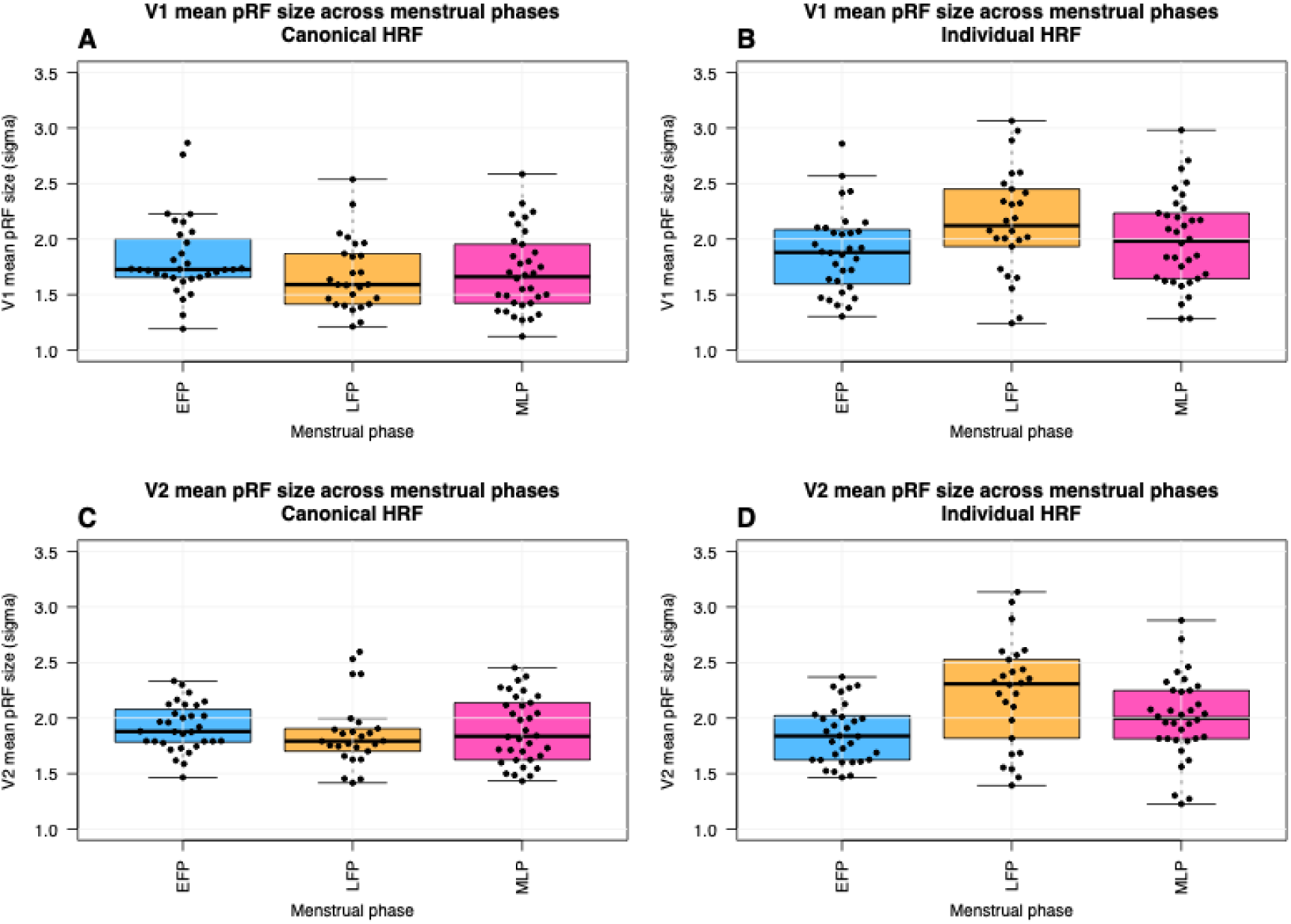
Boxplot illustrating average (mean) population receptive field size processed using a canonical (A and C) or individually mapped (B and D) haemodynamic response function (HRF), for cortical area V1 (A and B) and V2 (B and D). EFP= early follicular phase; LFP= late follicular phase; MLP= mid-luteal phase.

Statistical models suggest that neither oestradiol (*χ*^2^(1)=0.018) nor resProgesterone (*χ*^2^(1)=0.972) were associated with pRF size when modelled using a canonical HRF (both p>0.05). However, a significant association with oestradiol and slightly larger pRF sizes was discovered when using individually measured HRFs (*χ*^2^(1)=16.318; p<0.001; fixed effect estimate=0.0005°; 95% CIs=0.0003-0.0007; average variation attributed to oestradiol across a cycle in this sample=0.25°). In addition, resProgesterone accounted for a significant amount of pRF size variance when individually mapped HRFs were used for convolution, specifically with slightly narrower pRFs (*χ*^2^(1)=27.536; p<0.001; fixed effect estimate=-0.013°; 95% CIs=-0.018– -0.008; average variation attributed to resProgesterone across a cycle in this sample=0.20°). The results combined indicate that oestradiol increase the pRF sizes whereas resProgesterone reduces them.

## 4. Discussion

This study is the first to comprehensively investigate the influence of menstrual-related hormones on dynamic cerebrovascular physiology using advancing MRI methodology. We found that both oestradiol and additional progesterone variance were associated with an increase in hypercapnic CVR, both when quantified using perfusion and using % BOLD signal. Associations with oestradiol and resProgesterone were also found in HRF shape across the cortex. These changes were relatively small across the menstrual cycle in this sample but raise questions about relationships to menstrual symptomology and CVR changes in more major endocrine states, such as menopause.

This study extends previous work finding endocrine-sensitive changes in CVR across carotid and cerebral arteries (Debert et al., 2012; Krejza et al., 2013; Skinner et al., 2023) by demonstrating these increases across the cortex using both perfusion and % BOLD signal change. Similar to the endocrine influence on resting perfusion (Wright et al., 2026), we observed this effect globally, which aligns with the widespread distribution of endocrine receptors throughout the brain (B. McEwen, 2002; B. S. McEwen et al., 1997; Mosconi et al., 2024). Both oestradiol and progesterone have been shown to mediate endothelial nitric-oxide (NO) signalling (Chambliss & Shaul, 2002; Lekontseva et al., 2011; Mendelsohn & Karas, 1999; Miller, 1999; Pang et al., 2015; Yuan et al., 2016), which heavily influences vasodilation and thus CVR; elevated NO bioavailability may lead to greater vasodilatory capacity in response to CO_2_. Our findings of increased resting perfusion and increased vascular reactivity with ovarian hormones may also suggest that these hormones reduce vascular tone, which has been previously demonstrated (Sarrel, 1999; White, 2002), and therefore increases the vasodilatory reserve. Future work should further investigate the potential mechanisms behind this and the implications for symptomatology across the menstrual cycle and other endocrine events. For example, impaired CVR has been implicated in migraine (Dzator et al., 2021) and depressive mood (Lemke et al., 2010; Vakilian & Iranmanesh, 2010).

Additionally, hormone associations were observed in the visually-evoked HRF shape across visual cortical areas, which reflects the complex interplay between neural activity, neurovascular coupling, and CVR. Oestradiol was associated with an increase in peak amplitude (PA) and a faster time-to-peak (TTP). While it is not possible to disentangle the underlying mechanisms based on the HRF alone, it may be theorised that this association reflects the CVR relationship described above; an increase in blood flow and reactivity would lead to faster blood flow and greater vasodilation following neural activity. Amplified PA and shorter TTP are also independently associated, potentially reflecting the latency of neurovascular coupling processes (Thompson et al., 2014), which typically delays with age (West et al., 2019). Additional progesterone variance, however, was associated with a small increase in the FWHM, suggesting wider HRF shapes during high progesterone phases. This increased temporal duration of the HRF response may reflect a small delay in vasoconstrictive processes, lengthened neural activation, or disrupted neurovascular processes, and may be particularly influential for fMRI HRF convolution pipelines. This work extends findings from a previous study by Li *et al.,* (2025), which mapped cortical HRF across two different menstrual phases using resting-state fMRI estimation, rather than visually-evoked responses. While the use of two timepoints may have reduced the measured range of oestradiol, they also found that oestradiol was associated with HRF amplitude in a sample that reported menstrual migraines (a positive relationship was also found between progesterone and FWHM but this did not reach statistical significance in this cohort). This raises the possibility that such vascular fluctuations are mechanistically tied to menstrual symptoms, and a greater neuroendocrine effect may be observed in samples with severe reported menstrual symptoms.

One consequence of a menstrual endocrine influence on dynamic cerebrovascular functions is the interpretation of fMRI studies. Firstly, to make inferences about underlying neural activity between groups, BOLD fMRI studies must assume equal vascular contributions (e.g., Wright & Wise, 2018). However, this study suggests that this may not be the case if the groups of interest differ by menstrual phase or endocrine state. Secondly, BOLD fMRI analyses often involve convolution using a canonical HRF shape. However, if the underlying HRF shape differs by group, convolving all by the canonical shape may introduce noise. To address this, Experiment Two investigated the influence of using either a canonical or individually-mapped HRF shape during a pRF processing pipeline. We found that, while no associations between hormone and pRF size were apparent when using the canonical HRF, a significant effect was observed when individually mapped HRFs were used. This suggests that neural effects may be obscured if vaso-endocrine factors are not considered and emphasises the promise of using individually-mapped HRFs to reduce possible vascular variation.

After incorporating individually-mapped HRF convolutions, we found evidence that pRF sizes across V1 and V2 were positively associated with menstrual-related fluctuations in oestradiol and negatively associated with progesterone. The size of a pRF is theoretically modulated by the balance of excitatory and inhibitory processes that determine the underling receptive fields within the visual cortex, which provides a potential mechanism for this action. For example, oestradiol has been found to modulate intracortical excitability in the visual cortex across the menstrual cycle, with reduced paired-pulse suppression in high oestrogen phases (Schloemer et al., 2020). This change in pRF size may also have functional consequences, as pRF size has been associated with visual acuity (Silva et al., 2021). While visual sensitivity to achromatic stimuli and visual acuity appears to be relatively stable across the menstrual cycle, performance decreases in the luteal compared to the follicular phase have been reported using blue-yellow stimuli (short-wave automated perimetry; SWAP) (Y. Akar et al., 2005; Yucel et al., 2005), potentially reflecting greater endocrine sensitivity in the koniocellular pathway. Some menstruating people may also be more susceptible to such changes; luteal phase decreases in visual sensitivity was reported for both SWAP and achromatic stimuli in patients with migraine, rather than just SWAP stimuli (Yucel et al., 2005). Another study also found that while healthy controls did not demonstrate modulation in SWAP sensitivity, patients with diabetes did (M. Akar et al., 2005).

This study used advanced physiological MRI methodology to comprehensively image dynamic vascular function across the menstrual cycle. By testing three times across a cycle, we were able to potentially sample the full range of endocrine levels and detangle the contribution of oestradiol compared to progesterone. However, there are a number of limitations that should be considered. As reported in the previous resting study (Wright et al., 2026), the LFP timepoint was designed to sample the oestradiol ovulatory peak; however, this was missed in many participants, despite timing the phase days on previous literature (e.g., Draper et al., 2018), which may lead to an underestimation of oestradiol’s influence. Variation in oestrogen patterns within and between individuals is well documented (Celec et al., 2009; Gandara et al., 2007; Maman et al., 2023) and is especially important to consider when using methods like MRI that offer limited scheduling flexibility. This underscores the need to measure actual circulating hormone levels rather than relying on assumed menstrual phases, enabling direct comparison based on true hormone concentrations. The sample size for CVR analysis was also reduced compared to the full sample, limiting statistical power.

## 5. Conclusion

This study found evidence that dynamic cerebrovascular functions are sensitive to menstrual-related ovarian hormones. These effects were small but notable within the context of the regular menstrual cycle and raise questions about larger hormonal variations, such as within pregnancy or menopause. This may be a potential mechanism underlying menstrual symptomatology and has implications for fMRI studies that assume intact neurovascular coupling processes in women, regardless of menstrual staging, to make inferences about neural activity. As suggested by these retinotopy data, this added noise may be obscuring neuroendocrine-related effects.

## Acknowledgements

We would like to thank Wellcome Trust for their help in publication of this article. This research was funded in whole, or in part, by the Wellcome Trust (WT224267).

For the purpose of open access, the author has applied a CC BY public copyright license to any Author Accepted Manuscript version arising from this submission.

## Data statement

### Credit statement

MEW: Conceptualization, Methodology, Investigation, Formal analysis, Writing – original draft, Writing – review & editing

ID: Formal analysis, Writing – review & editing

CC: Methodology, Investigation, Formal analysis, Writing – review & editing

SD: Project Administration, Investigation, Writing – review & editing

JJS: Conceptualization, Methodology

DDS: Software, Writing – review & editing

KM: Conceptualization, Methodology, Formal analysis, Writing – original draft, Writing – review & editing, Supervision, Funding acquisition.

### Conflicts of interest

The authors have no conflicts of interests to declare.

## Notes

### Competing Interest Statement

The authors have declared no competing interest.

